# Identifying Molecular Determinants and Therapeutic Targets in Luminal B Breast Cancer: A Systems Biology Approach

**DOI:** 10.1101/2025.05.25.656027

**Authors:** Yousef Saeidi, Masoud Ghorbani, Ali Najafi, Mehrdad Moosazadeh Moghaddam

## Abstract

Luminal B breast cancer (LBBC) is on the rise worldwide, with both incidence and mortality rates steadily increasing. Early detection proves difficult due to its aggressive characteristics, most notably its heightened proliferation rate and the complex interplay of molecular biomarkers, particularly in more advanced stages. In the present study, we conducted an *in silico* analysis of LBBC cell lines using the Gene Expression Omnibus (GEO) microarray dataset, which includes 30 LBBC and 11 normal samples. Differentially expressed genes (DEGs) were identified using RStudio. A series of analyses, including cancer data interrogation via pan-cancer analysis, eXpression2Kinases (X2K), and the Cancer Dependency Map (DepMAP), was carried out to elucidate the underlying signaling pathways, transcription factors (TFs), and kinases, as well as to perform stemformatics analysis. Protein–protein interaction (PPI) networks were reconstructed using STRING and Cytoscape, enabling the identification of co-expressed and hub genes through the cytoHubba plug-in. Of note, FGF2, EGFR, ADIPOQ, LIPE, MET, IGF1, FGF1, EGF, FGF7, and PPARG were identified as the top 10 driver genes; RELA, PPARG, CTCF, EGR1, and NFE2L2 were detected as predominant TFs in LBBC; and CDK1, CDK2, MAPK3, CSNK2A1, and MAPK14 were identified as potential biomarkers through hub gene clustering. Further analysis indicated hsa-mir-221-3p and hsa-mir-29a-3p as key miRNAs targeting LBBC-related genes. Collectively, our findings highlighted LBBC-associated genes, TFs, miRNAs, and pathways as prospective biomarkers, providing insights into LBBC diagnosis and therapeutic approaches.

## Introduction

Breast cancer is the second most commonly diagnosed cancer worldwide and the leading cause of cancer-related mortality among women[1]. According to the PAM50 gene expression profiling algorithm, breast cancer is classified into five intrinsic molecular subtypes: Luminal A (LumA), Luminal B (LumB), HER2-enriched (HER2-E), Basal-like, and Normal-like[2]. Among these subtypes, LumB breast cancer is considered as a clinically aggressive subtype, accounting for approximately 15–20% of all breast cancer cases[2, 3]. LumB tumors are characterized by the expression of hormone receptors (estrogen and/or progesterone receptors), and distinguished by higher proliferation rates and elevated Ki-67 expression (a marker of cellular proliferation), compared with LumA tumors [3]. This phenotype is associated with poorer clinical outcomes and reduced responsiveness to standard endocrine therapies, highlighting the need for novel therapeutic approaches [4, 5].

Advancements in high-throughput sequencing and omics technologies, such as genomics, transcriptomics, proteomics and metabolomics, have revolutionized cancer research by unveiling the molecular complexity of tumors [6, 7]. These technologies have the ability to identify critical biomarkers for early detection, cancer stratification and personalized treatment approaches [8–10]. In the context of LumB breast cancer, integrating multi-omics data provides a holistic view of tumor biology, revealing distinct molecular landscapes that drive tumor aggressiveness[11]. Previous studies have elucidated specific genetic alterations in LumB tumors, notably TP53 mutations and heterogeneity in HER2 expression[12, 13]. Additionally, transcriptomic analyses have provided insights into the roles of transcription factors, kinases and non-coding RNAs (ncRNAs), including microRNAs (miRNAs), in the pathogenesis of LumB breast cancer [14, 15].

Furthermore, cancer-associated fibroblasts (CAFs) within the tumor microenvironment (TME) are implicated in LumB tumor progression, therapy resistance and metastasis by modulating signaling pathways critical to proliferation, migration and angiogenesis[16–18]. Despite these advances, many findings are fragmented and a systematic understanding of the molecular regulators driving LumB aggressiveness remains elusive.

To address these gaps, integrative bioinformatics approaches have emerged as powerful tools to synthesize diverse omics data. Protein-protein interaction (PPI) networks, transcriptional regulatory networks and epigenetic modifications provide insights into the interplay between genetic and non-genetic factors in tumor biology. Additionally, tools, such as Cancer Dependency Map (DepMap), and miRNA network analyses have advanced our understanding of tumor vulnerabilities, revealing potential therapeutic targets with high precision[19, 20].

This study employed a multi-omics framework to investigate LumB breast cancer at the molecular level. Using transcriptomic, epigenomic, and regulatory network analyses, we aimed to identify key molecular features and therapeutic targets that define LumB tumor biology. By integrating data from PPI networks, signaling pathway enrichment, cancer dependency maps, and miRNA-circRNA interactions, this study sought to uncover novel insights into tumor progression and therapeutic resistance. The findings had the potential to enhance diagnostic precision, stratify patients based on molecular characteristics, and inform the development of personalized therapeutic strategies, ultimately improving clinical outcomes for LumB breast cancer patients.

## Materials and Methods

### Gene Expression Analysis

Gene Expression Omnibus (GEO), a database for gene expression profiling and RNA methylation profiling maintained by the National Center for Biotechnology Information (NCBI), not only adheres to community-driven reporting standards, but also ensures the inclusion of several key study components such as raw data, processed data and descriptive metadata[21]. Data for whole-transcriptome expression analysis of luminal B breast cancer (LBBC) were obtained from the NCBI GEO database (www.ncbi.nlm.nih.gov/geo). The gene expression profile dataset, with accession number GSE45827, was retrieved from GEO (platform: GPL570 [HG-U133_Plus_2] Affymetrix Human Genome U133 Plus 2.0 Array), comprising 30 LBBC and 11 normal samples. Data expressed in this study were analyzed by R-studio. P-values < 0.05 was considered to be statistically significant.

### Protein-Protein Interaction (PPI) Network Analysis

The STRING database integrates all known and predicted associations among proteins, including both physical interactions and functional associations, through collecting and evaluating evidence from several sources [20]. STRING is also a search tool for retrieval of interacting gene databases (www.string-db.org) which integrates both known and predicted protein-protein interactions (PPIs) that are used to predict functional interactions between DEGs (high confidence score 0.700 was set as the cut-off criteria to construct PPI network). Furthermore, the CytoHubba (version 0.1) plugin of the Cytoscape software (version 3.10.2; www.cytoscape.org) was used to identify important genes.

### Enrichment Analysis

WebGestalt 2024 marks a significant upgrade of the functional enrichment analysis platform (www.webgestalt.org). This update not only brings the database up to date but also enhances the tool’s capabilities by incorporating support for metabolomics and introducing new pathways, networks and gene signatures[22]. A P-value of equal to/fewer than 0.05 was a significant boarder in gene ontology (GO) terms and pathways.

### Cancer Dependency Map (DepMAP) Analysis

The DepMap database (www.depmap.org), building off of the original Cancer Cell Line Encyclopedia (CCLE) project, creates data and tools that can be used and shared by researchers [19]. DepMap aids researchers in identifying genes that are crucial for the survival of cancer cells (genes with negative effects) and can be proposed as potential therapeutic targets. For example, genes with an effect score of less than -0.5 may represent promising candidates for the development of new therapeutic agents.

### Cancer Data Analysis

The University of ALabama at Birmingham CANcer data analysis Portal (UALCAN) (www.ualcan.path.uab.edu) is an interactive web resource for analyzing cancer omics data. The UALCAN web portal provides access to publicly available cancer transcriptome data, primarily from The Cancer Genome Atlas (TCGA), allowing researchers to explore gene expression and its association with patient survival, tumor stage and other clinical features across multiple cancer types. The platform offers customizable plots and statistical analyses, facilitating the identification of potential biomarkers or therapeutic targets[23].

### Stemformatics Analysis

Stemformatics (www.stemformatics.org) is an online platform designed to provide high-quality visualization and analysis of stem cell-related gene expression data. It hosts a curated collection of publicly available datasets from various platforms, including microarray and RNA-seq, with a focus on stem cell biology and differentiation processes. Stemformatics is particularly useful for bioinformatics-driven insights into stem cell biology, facilitating hypothesis generation and validation in regenerative medicine and developmental biology[24].

### Detection of Transcription Factors (TFs) and Kinases

Transcription Factors (TFs) potentially regulating LBBC-related genes were identified using the ChIP Enrichment Analysis (ChEA) database, which provides data on eukaryotic TFs, binding motifs, experimentally validated binding regions and target genes[25]. Moreover, eXpression2Kinases (X2K) was used to rank putative TFs, protein complexes and kinases likely driving the observed transcriptomic changes in LBBC.

### Small RNA Analysis

miRNet (www.mirnet.ca) is a comprehensive web-based platform designed to facilitate the integrative analysis of microRNA (miRNA)-centric regulatory networks. The database aggregates a wide range of experimentally validated and predicted miRNA-target interactions, including miRNA-gene, miRNA-protein, miRNA-disease, etc. Top miRNAs targeting LBBC-related genes were selected and ranked based on P-value (P ≤ 0.05)[26].

## Results

### Identification of Differentially Expressed Genes (DEGs)

The genes with different expressions were screened among the defined groups (30 LBBC and 11 normal samples). The Limma R package was used to identify DEGs. A P-value < 0.05 and |LogFC| > 2.0 were considered to be statistically significant. According to the statistics, a volcano plot is a scatter plot used to quickly spot changes in big data sets consisting of replicate data. Significance and fold-change are plotted on the y and x axes, respectively. Points of interest that display both large amplitude fold-changes (x axis) and high statistical significance are indicated by -log10 of p value, y axis. Those points with a fold-change less than two (log2 < 2) are represented in gray on this graph (Figure 1a).

**Figure 1.**
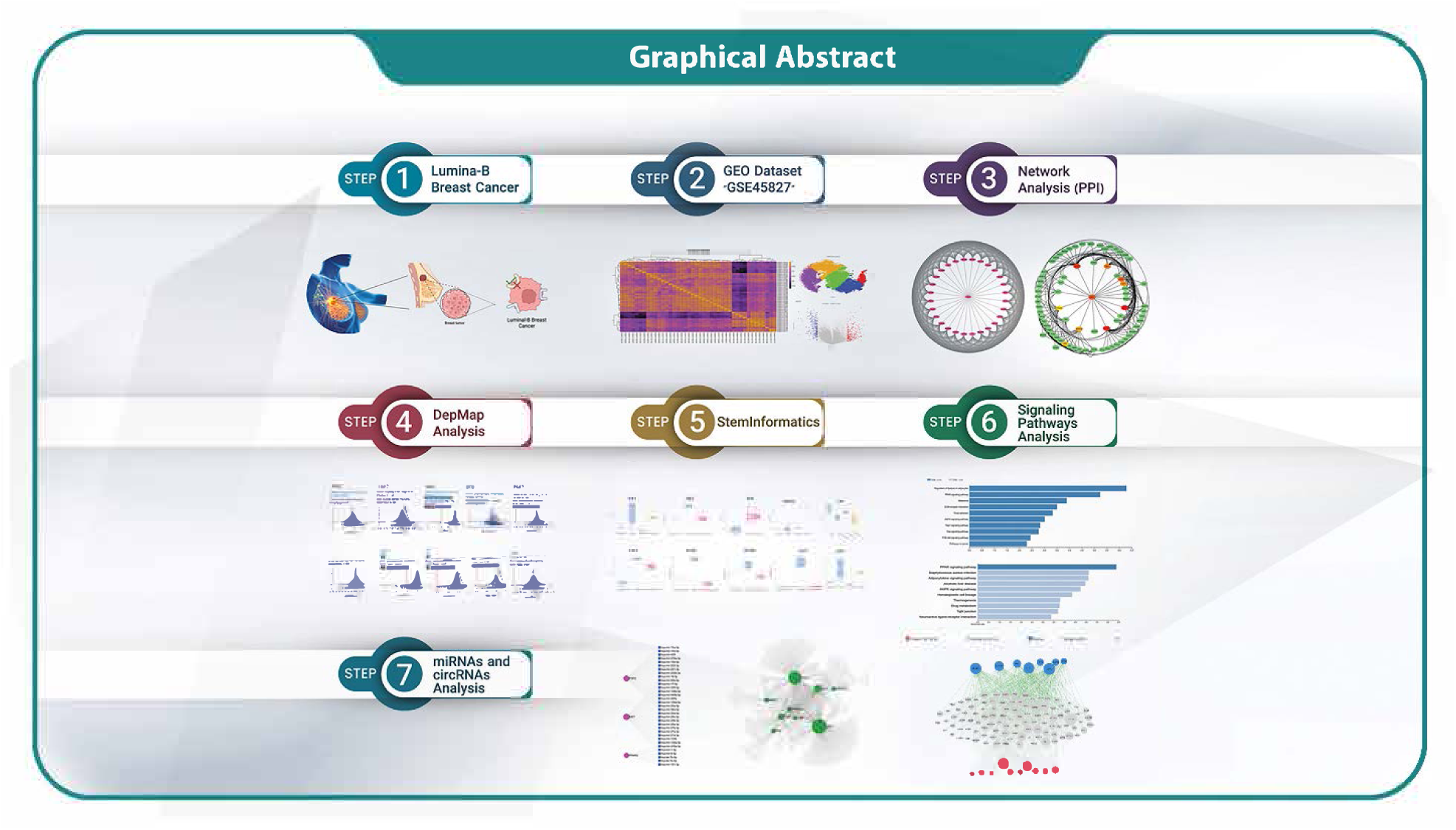
Graphical Abstract of the Integrated Multi-Omics Workflow for Luminal-B Breast Cancer Analysis.

The interactions of up and down-regulated genes were investigated by using the STRING database. Cytoscape software v 3.10.2 (cytoHubba plugin) analysis was carried out to identify hub genes (Figure 1b). The hub gene list was compiled by analyzing gene expression matrix data and drawing co-expression correlation coefficient heatmap. The co-expression study revealed the relationship among LBBC-associated genes.

### Pan cancer Analysis

The methylation analysis of hub genes was conducted using the web-based UALCAN platform, identifying hub genes, followed by the construction of their network(Figure 2a, b). Boxplots indicates inverse relation between promoter methylation status and gene expression profile of LBBC in TCGA invasive breast cancer. Our results showed that the expression of genes in the data matrix promoter levels of methylation in 6 up genes, including fibroblast growth factor 1

**Figure 2:**
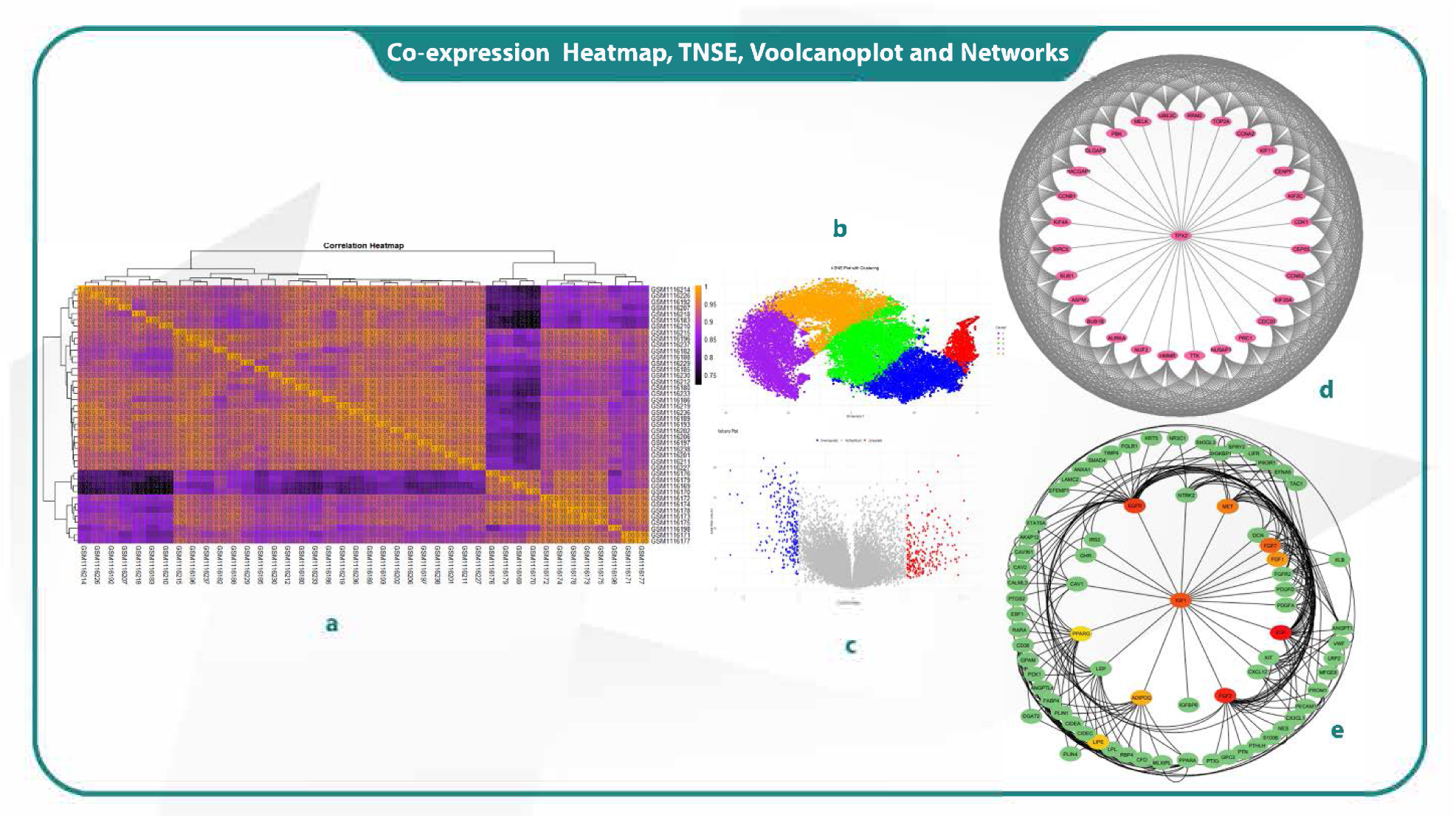
Co-expression heatmap analysis (a), Sample clustering based on t-SNE (b), Volcano plot of expression changes (c), Network of downregulated genes (d), Network of upregulated genes (e).

(FGF1), proto-oncogene, receptor tyrosine kinase (MET), insulin like growth factor 1 (IGF1), peroxisome proliferator activated receptor gamma (PPARG), lipase E, hormone sensitive type (LIPE), epidermal growth factor receptor (EGFR) and 6 down genes, including cyclin dependent kinase 1 (CDK1), kinesin family member 11 (KIF11), hyaluronan mediated motility receptor (HMMR), PDZ binding kinase (PBK), cyclin A2 (CCNA2) and components of NDC80 kinetochore complex (NUF2) were found to be driver genes (Figure 2).

### Analysis of Cancer-Related Gene Dependencies Using DepMap Data

This study analyzed the dependency of various cancer-related genes across multiple cell lines using data from the DepMap database. CRISPR-Cas9 and RNAi screening were employed to assess the essentiality of genes for cell survival. Each gene’s dependency was quantified using gene effect scores, with negative scores indicating higher dependency. Key findings included genes like EGFR and MET, which were classified as highly selective and essential in a significant number of cancer cell lines, suggesting their potential as therapeutic targets. In contrast, genes like FGF1, FGF2 and EGF showed limited essentiality, indicating their less importance across these cell lines. PPARG and LIPE showed a mix of CRISPR and RNAi data, with some selectivity but not as broadly essential as EGFR (Figure 3).

**Figure 3:**
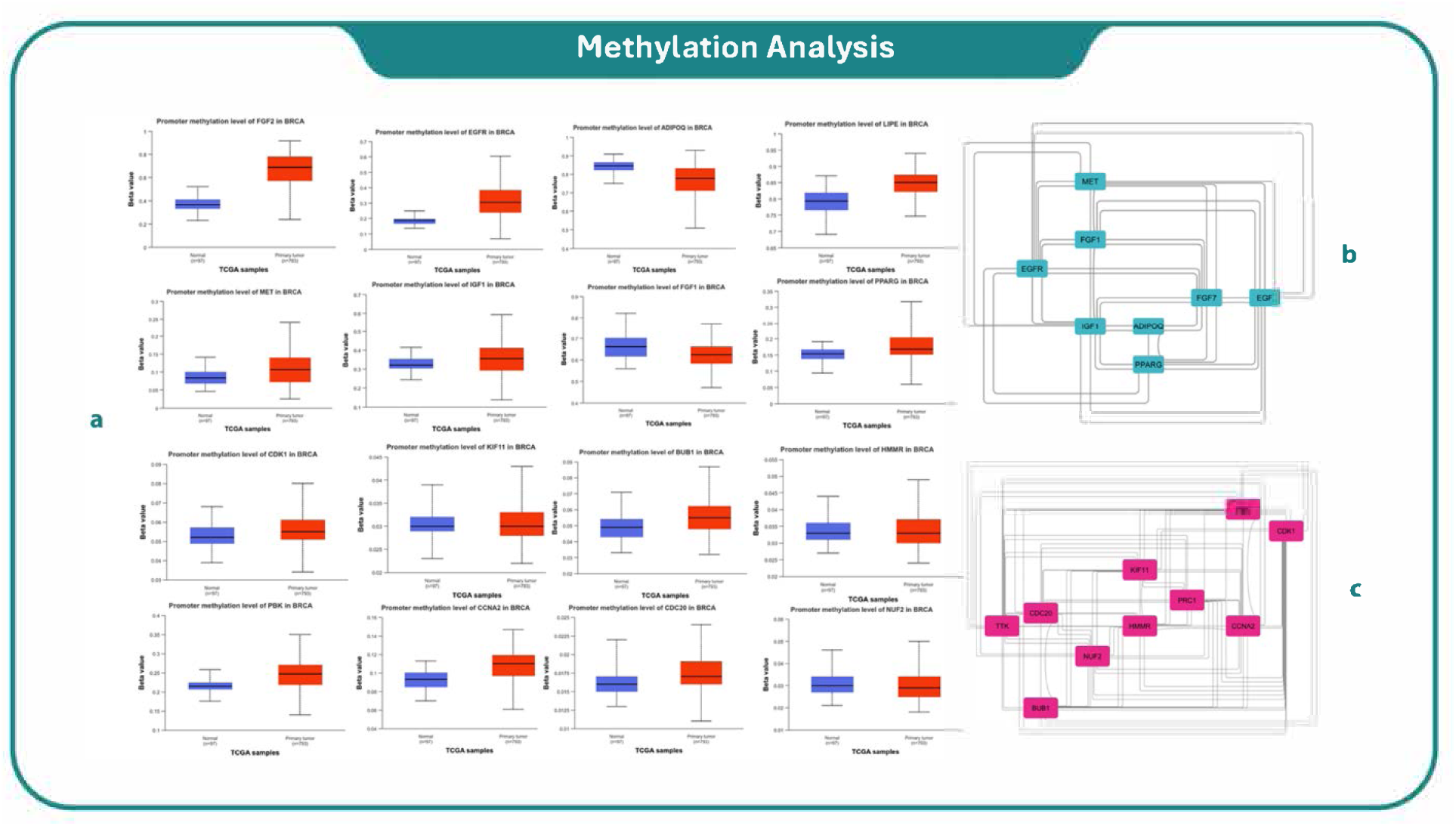
Methylation analysis to find driver genes (a), Up and down driver gene networks (b and c)

### Stemformatics Analysis

Stemformatics analysis was conducted to investigate the role of specific genes in cancer stem cells. Our analysis revealed the pivotal roles of EGFR, IGF1, PPARG, EGF, LIPE, ADIPOQ, MET, FGF1, and FGF7 in the elevated expression seen in fibroblasts. These genes are crucial in modulating key pathways that influence fibroblast activity, particularly in the tumor microenvironment. Their upregulation suggests a significant contribution to fibroblast-cancer cell interactions, promoting processes such as cell proliferation, migration and angiogenesis. These findings underscored the importance of fibroblast-related gene expression in cancer progression, highlighting potential therapeutic targets within the tumor stroma (Figure 4).

**Figure 4:**
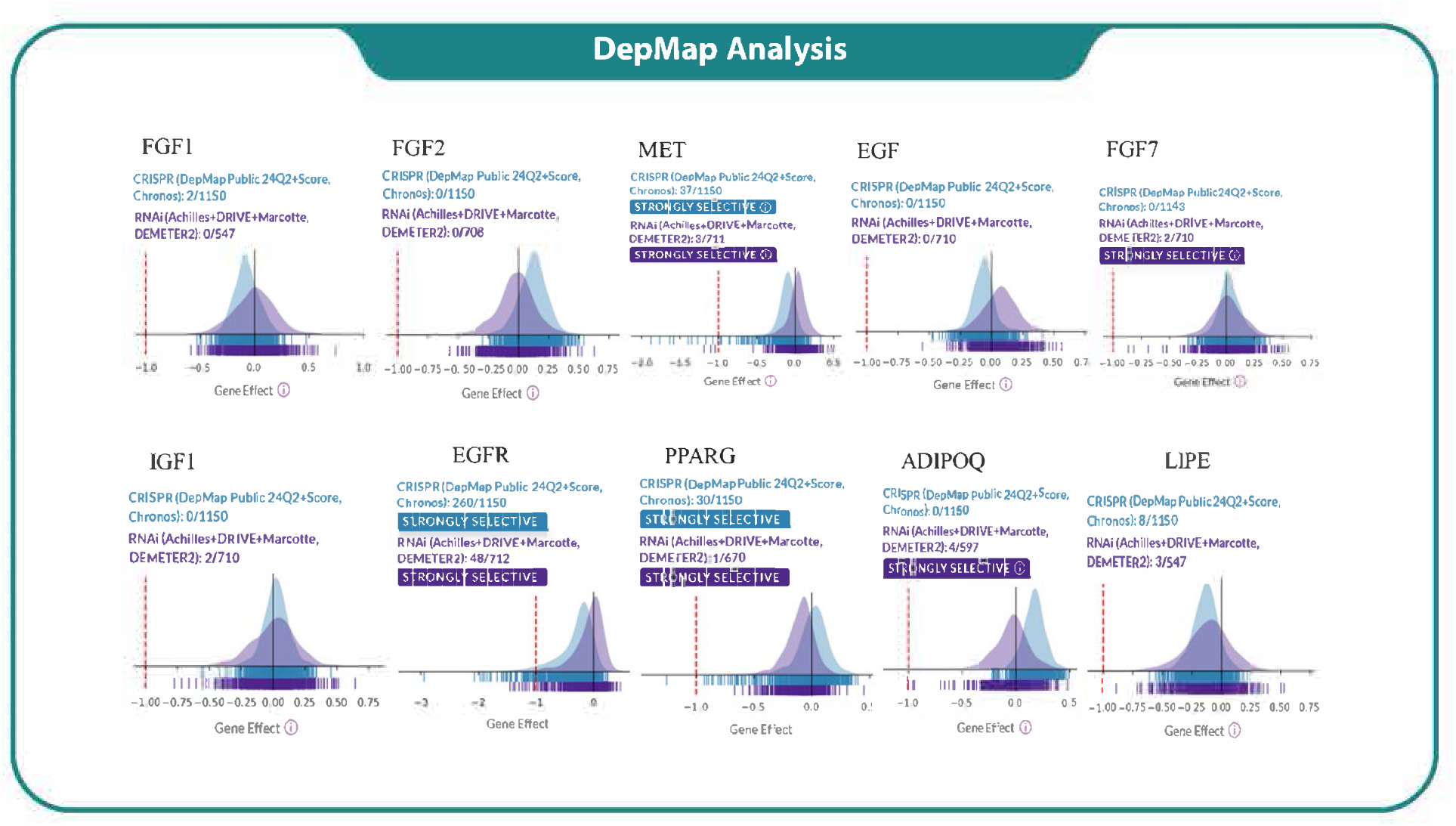
DepMap analysis of the dependency of tumor cell line panels in CRISPR (blue) and RNAi (violet) databases on the indicated driver genes.

### Signaling Pathway Analysis

This study identified several keys signaling pathways significantly enriched in our dataset through the analysis of gene expression data. Regulation of Lipolysis in Adipocytes was found to be the most prominent pathway, showing the highest degree of enrichment. Moreover, the PPAR signaling pathway was notably enriched, suggesting its potential role in the regulation of metabolic processes linked to cancer progression. Further pathways of interest included the Melanoma pathway and the ECM-Receptor Interaction. The Focal Adhesion pathway, which is crucial for cell migration, invasion and processes that are often dysregulated in cancer cells, was found another significant finding. We also identified several oncogenic signaling cascades, such as PI3K- Akt, Ras, and Rap1 signaling pathways, all of which are frequently implicated in cancer development, cell proliferation and survival. Moreover, the AMPK signaling pathway, which plays a key role in cellular energy homeostasis, was highlighted as a significant pathway, potentially linking metabolic stress responses to cancer progression (Figure 5 a, b).

**Figure 5:**
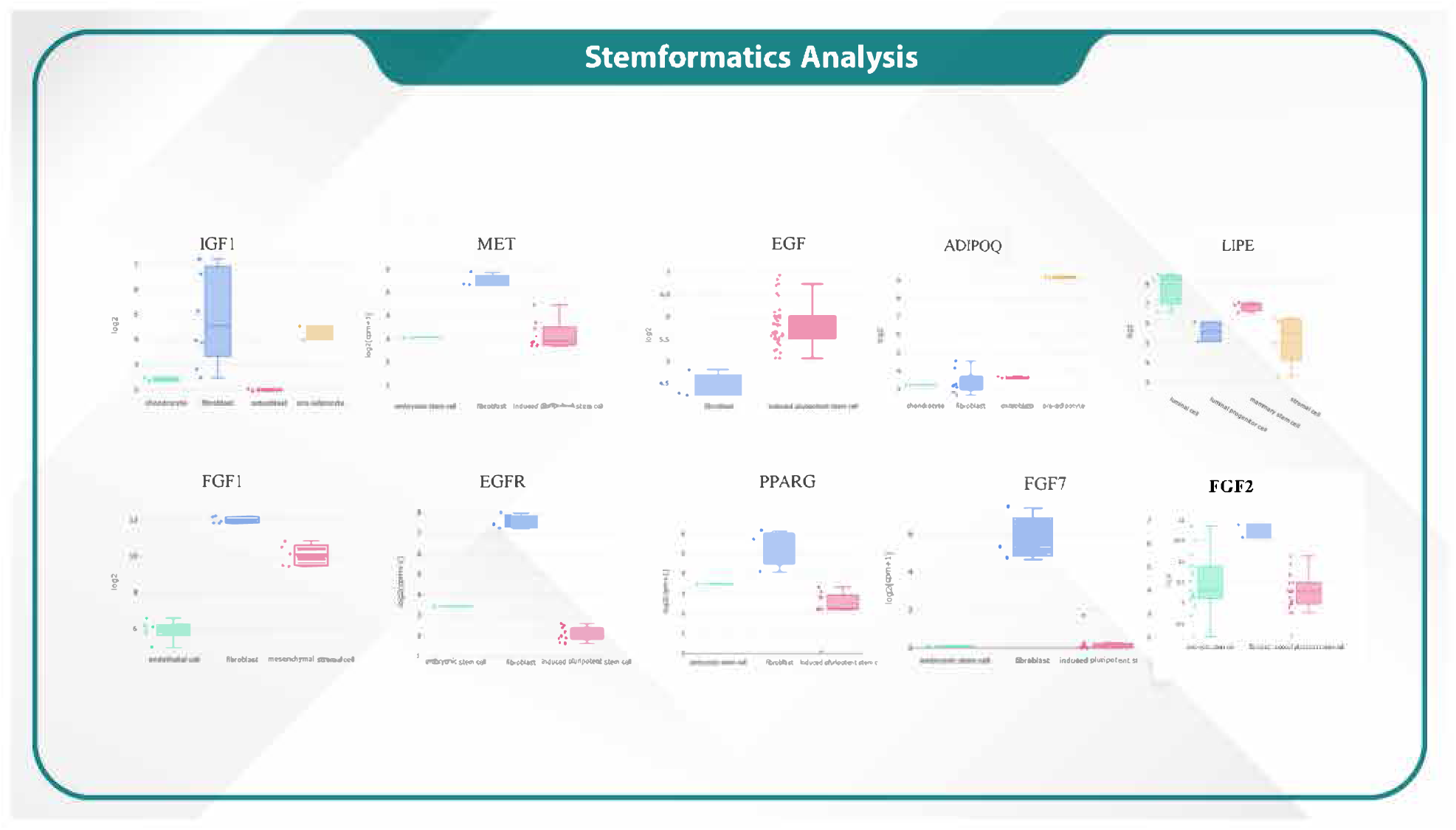
Stemformatic analysis to demonstrate increased expression of driver genes in cancer stem cells

### Stemformatics Analysis

Gene expression analysis using the Stemformatics database revealed distinct expression patterns across different cell types. Notably, the fibroblast cells exhibited the highest expression levels for several key genes. Specifically, IGF1, MET, EGF, EGFR, PPARG and FGF7 showed significantly elevated expression in fibroblasts compared with other cell types, indicating a prominent role of these genes in fibroblast functions. The findings suggest that fibroblasts have a unique transcriptomic profile, in which these genes are upregulated, potentially implicating their importance in cellular differentiation and tissue repair processes (Figure 6).

**Figure 6:**
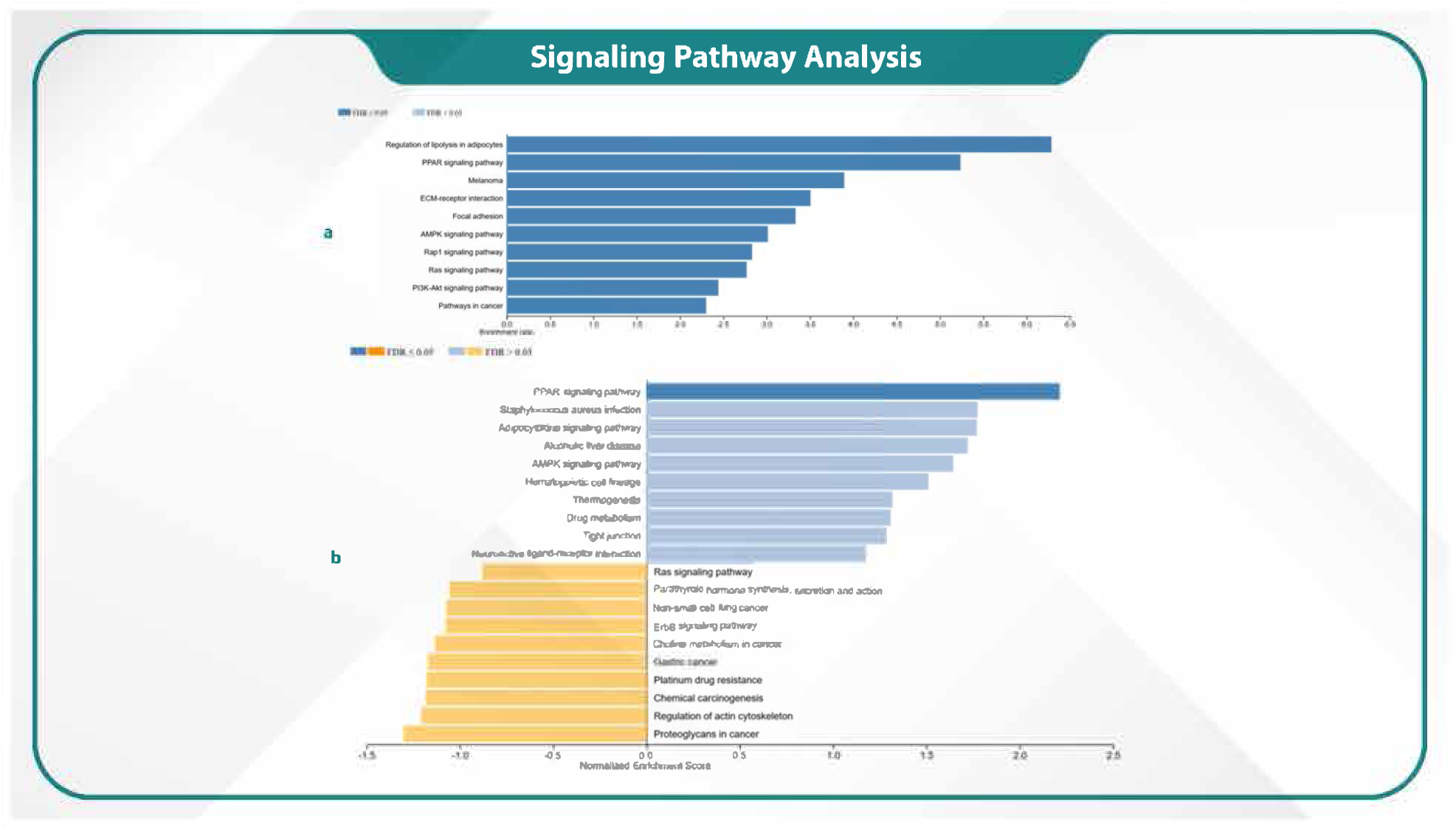
Signaling pathways analysis of up- and down-regulated genes (a, b)

### Identification of kinases and transcription factors (TFs)

X2K was used to identify the key TFs, kinases, and intermediary proteins involved in the regulation of gene expression. Our results revealed that RELA, PPARG, EGR1, NFE2L2, and TP63 were the most significant TFs targeting the greatest number of genes associated with LBBC. Among 10 TFs, RELA and PPARG showed the most interactions with intermediate proteins and kinases (Fig. 7).

**Figure 7:**
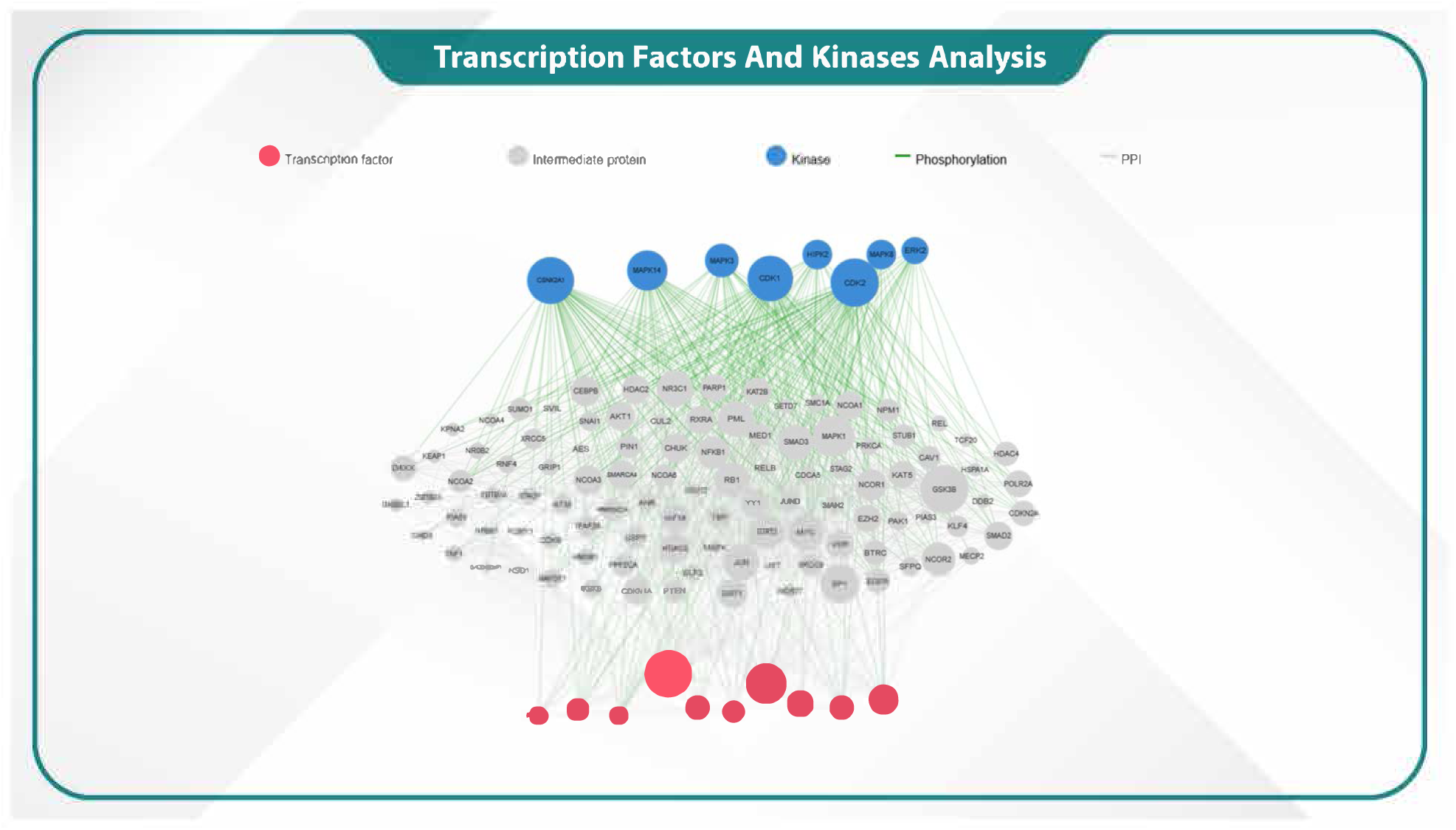
The interaction of transcription factors (TFs; red spots) and kinases (blue spots) with hub genes.

### miRNA Network Analysis

We identified microRNA (miRNA) networks for the genes FGF7, FGF2, IGF1, ADIPOQ, EGF, FGF1, LIPE, MET and PPARG using the miRNet database. In addition, miRNA networks were detected for FGF2, MET and PPARG, indicating potential regulatory interactions involving these miRNAs. These findings highlighted the intricate post-transcriptional regulation of these genes through both microRNAs (Figure 8).

**Figure 8:**
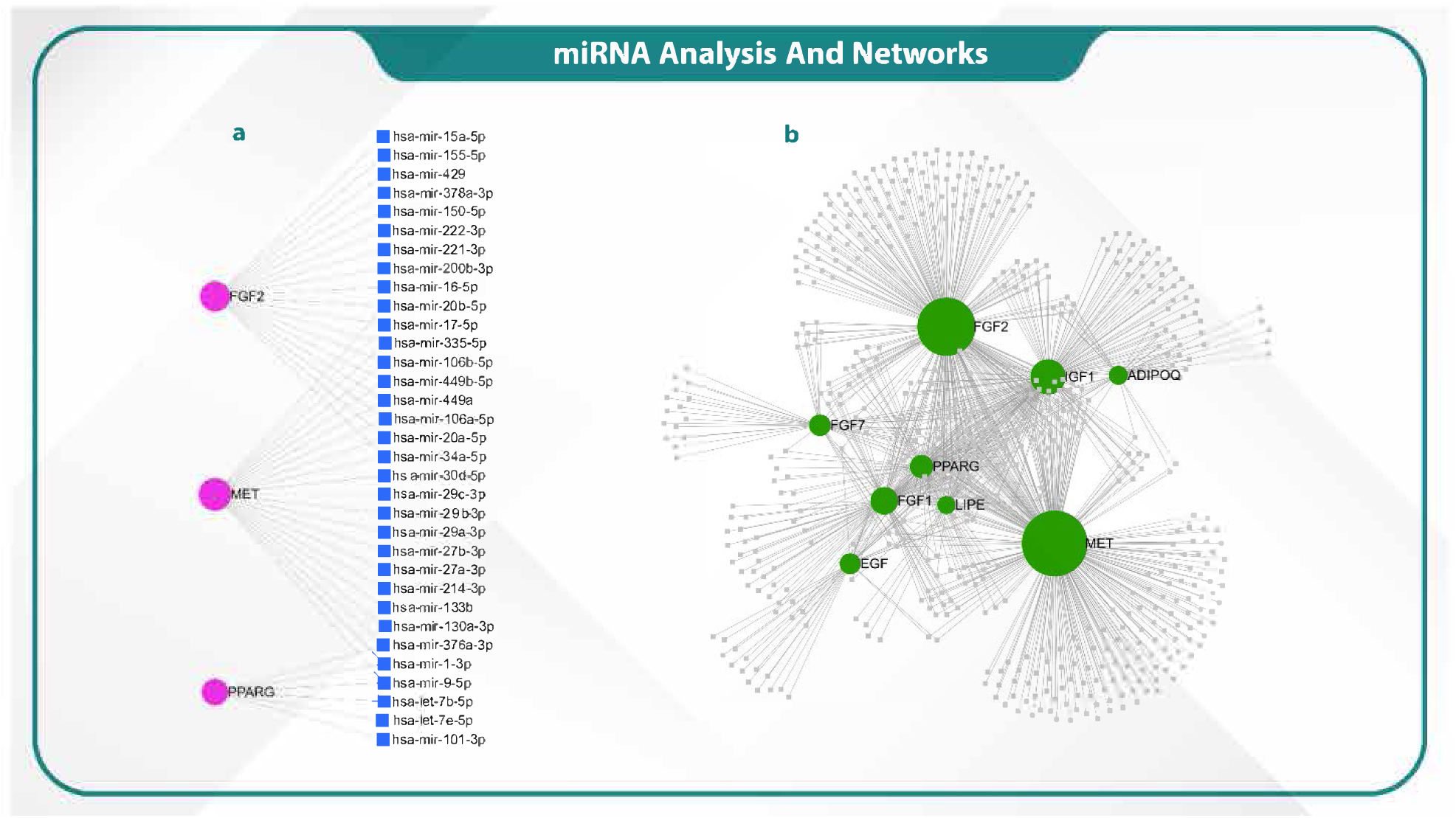
Identification of the key miRNAs and genes involved in LumB.

## Discussion

The present study could successfully identify the main molecular regulators of LumB breast cancer (LBBC) through transcriptomic analysis, opening a potential window for new therapeutic targets. Indeed, our analysis focused specifically on key entities, including hub genes, TFs, miRNAs, kinases, CAFs and PPIs.

The dataset (GSE45827) analyzed in our study contained 2,206 downregulated and 945 upregulated genes (from 11 normal and 30 tumor samples). The DEG analysis revealed 18 hub genes with significant prognostic potential, among which 10 were upregulated genes (including FGF2, EGFR, ADIPOQ, LIPE, MET, IGF1, FGF1, EGF, FGF7 and PPARG but 8 were downregulated genes (including CDK1, KIF11, BUB1, HMMR, PBK, CCNA2, CDC20 and NUF2. The discovery of such genes, despite significant advances in understanding the molecular causes of LumB breast cancer, may help identify potential targets for the development of novel therapeutic agents.

There are studies showing altered methylation patterns of several hub genes in cancer, reflecting their key roles in tumor growth and survival. FGF1 and FGF2 are involved in cell proliferation and angiogenesis [27], while EGFR regulates crucial signaling pathways such as MAPK and PI3K- Akt, making it a promising therapeutic target [28, 29]. In addition, MET was demonstrated to play a role in metastasis and survival, with its methylation changes suggesting disrupted regulation [30].

The upregulation of metabolic genes, such as ADIPOQ, LIPE, IGF1 and PPARG, highlights the tumor’s metabolic reprogramming to drive in nutrient-rich environments [31]. Our results showed that the epigenetically-modified methylation can serve as a dual regulator so as to not only facilitates the DNA self-assembly process but can also be used as a universal biomarker for cancer [32–35]. Moreover, our investigation of the strongly selective dependencies identified for MET, EGFR, and PPARG reinforce their established roles as oncogenes or key modulators in cancer biology[34].

Results from deep map highlights the importance of integrating CRISPR and RNAi datasets for a comprehensive understanding of gene essentiality in cancer [36]. Our results showed the dependency patterns of FGF7, ADIPOQ and LIPE underscore emerging vulnerabilities that may be context-specific, offering potential new therapeutic avenue[37, 38]. Additionally, the lack of dependency for genes such as FGF1, FGF2 and EGF suggests functional redundancy or limited cancer-specific roles, emphasizing the necessity of multi-gene pathway analysis.

Stemformatics analysis revealed the elevated expression of MET, EGFR, IGF, FGF1 and FGF7 highlights their importance in CAFs are the principal population of stromal cells in lumB tumors[39–41]. Our results demonstrated that PPARG and ADIPOQ, whose central roles were previously discovered in CAFs, also exhibit increased expression in fibroblasts.

Because EGF has the highest expression level in iPSCs, the identification of such potential genes may help develop novel therapeutic strategies.

According to this study, Rap1 and Ras pathways are essential for controlling cell adhesion and migration, and they can affect these functions by sharing adhesion molecules or downstream kinases. Interactions between Rap1 and Ras may have a synergistic effect on cell invasion[42, 43]. Simultaneous targeting of Rap1 and Ras pathways can serve as an effective strategy to decrease cancer cell invasion and metastasis.

Importantly, studies focusing on such transcription factors and kinases may help identify downstream Rap1 and Ras signaling pathways including transcription factors (TP53, RELA, PPARG) and kinases (CDK1, CDK2, MAPK14) [44]. These kinases, which phosphorylate TFs, are essential for stress responses and cell cycle regulation.

Lastly, the intricate regulatory network involving miRNAs emphasizes their roles in modulating key genes such as MET, FGF2, and PPARG. Our research showed that hsa-mir-221-3p and hsa-mir-29a-3p play crucial roles in tumorigenesis and angiogenesis [45, 46]. Such miRNAs target critical components of signaling pathways involved in cancer, metabolism and cellular differentiation.

In conclusion, findings from this study shed light on the critical factors associated with the development of lumB, paving the way for improved therapeutic interventions. Of great note, integrating transcriptomics data can result in developing novel treatment approaches for patients with breast cancer.

## Declarations

### Ethical Approval and Consent to participate

The datasets analyzed in this study were obtained from [NCBI repository GSE45827], which provides de-identified human data. Since the data were publicly available and anonymized, no additional ethical approval or consent was required for this secondary analysis.

## Data Availability

The datasets generated during and/or analysed during the current study are available in the GEO DataSets accession number [GSE45827], https://www.ncbi.nlm.nih.gov/geo/query/acc.cgi?acc=GSE45827

## Contributions

**Yousef Saeidi:** Methodology, Data curation, Writing-original draft.

**Masoud Ghorbani:** Investigation, Data curation.

**Ali Najafi:** Validation, Software, Data curation.

**Mehrdad Moosazadeh Moghaddam:** Writing-review & editing, Supervision, Project administration.

## Acknowledgements

The authors would like to acknowledge the reviewers for their helpful and constructive comments on this manuscript.

